# Toward Broad Spectrum DHFR inhibitors Targeting Trimethoprim Resistant Enzymes Identified in Clinical Isolates of Methicillin-Resistant *Staphylococcus aureus*

**DOI:** 10.1101/648808

**Authors:** Stephanie M. Reeve, Debjani Si, Jolanta Krucinska, Yongzhao Yan, Kishore Viswanathan, Siyu Wang, Graham T. Holt, Marcel S. Frenkel, Adegoke A. Ojewole, Alexavier Estrada, Sherry S. Agabiti, Jeremy B. Alverson, Nathan D. Gibson, Nigel D. Priestly, Andrew J. Wiemer, Bruce R. Donald, Dennis L. Wright

## Abstract

The spread of plasmid borne resistance enzymes in clinical *Staphylococcus aureus* isolates is rendering trimethoprim and iclaprim, both inhibitors of dihydrofolate reductase (DHFR), ineffective. Continued exploitation of these targets will require compounds that can broadly inhibit these resistance-confering isoforms. Using a structure-based approach, we have developed a novel class of ionized non-classical antifolates (INCAs) that capture the molecular interactions that have been exclusive to classical antifolates. These modifications allow for a greatly expanded spectrum of activity across these pathogenic DHFR isoforms, while maintaining the ability to penetrate the bacterial cell wall. Using biochemical, structural and computational methods, we are able to optimize these inhibitors to the conserved active sites of the endogenous and trimethoprim resistant DHFR enzymes. Here, we report a series of INCA compounds that exhibit low nanomolar enzymatic activity and potent cellular activity with human selectivity against a panel of clinically relevant TMP^R^ MRSA isolates.

---

Antibacterial resistance is a growing healthcare and public health crisis worldwide. The rapid dissemination of antibiotic resistance has diminished the efficacy of many once reliable therapeutics. In fact, resistance to every class of antibiotics has been observed clinically. The Review on Antimicrobial Resistance projected that drug resistant infections will be responsible for more than 10 million deaths a year by 2050 and cost the global economy over 100 trillion USD. Among the most prevalent pathogens that have been identified as particular concern are methicillin and vancomycin-resistant strains of *Staphylococcus aureus*^1^.

Methicillin resistant *Staphylococcus aureus* (MRSA), an opportunistic gram-positive bacterium, is the leading cause of healthcare associated infections as well as invasive systemic infections, pneumonia and skin and soft tissue infections (SSTIs) worldwide. The CDC reports over 80,000 invasive MRSA infections annually in the United States, more than 11,000 of which are fatal, which has prompted the CDC to classify drug resistant MRSA as a ‘Serious Threat’^2^.

The antifolate combination of trimethoprim and sulfamethoxazole (co-trimethoxazole), marketed as Bactrim or Septra, is a first line treatment for community acquired skin and soft tissue MRSA infections. Trimethoprim targets dihydrofolate reductase (DHFR) which is responsible for the NADPH-dependent reduction of dihydrofolate (DHF) to tetrahydrofolate (THF). DHFR is the only source for the recycling of THF in the cell. When employed in conjunction with sulfamethoxazole, which targets dihydropteroate synthase, this powerful synergistic antibacterial combination results with potent coverage against both Gram-negative and Gram-positive pathogens. Due to its broad spectrum of activity, oral bioavailability and general tolerability, prescriptions of TMP-SMX numbered more than 21 million in 2013, putting it in the group of top ten oral antibiotics prescribed^3^.

Currently, trimethoprim is the sole FDA-approved antibiotic targeting DHFR. A second compound, iclaprim, a structurally similar DHFR inhibitor with anti-staphylococcal activity, has recently completed a Phase III clinical trial for acute bacterial skin and skin structure (ABSSI) infections^4^. DHFR inhibitors are historically grouped into two classes: lipophilic and classical. Trimethoprim and iclaprim are lipophilic antifolates as they contain a 2,4-diaminopyrimidine pharmacophore and passively diffuse into the cytosolic space. Methotrexate and pemetrexed, both chemotherapeutics, are known as classical antifolates as they possess a glutamate moiety in their structure, Figure 1. As mimics of the natural substrate DHF, classical antifolates show high affinity to all DHFR enzymes, however due to the negatively charged glutamate tail (net charge= −2), these compounds must be actively transported into the cell via specific folate carriers. Since bacteria do not have these transport mechanisms, classical antifolates do not show significant antibacterial efficacy despite powerful inhibition of bacterial DHFR.

**Figure 1:**
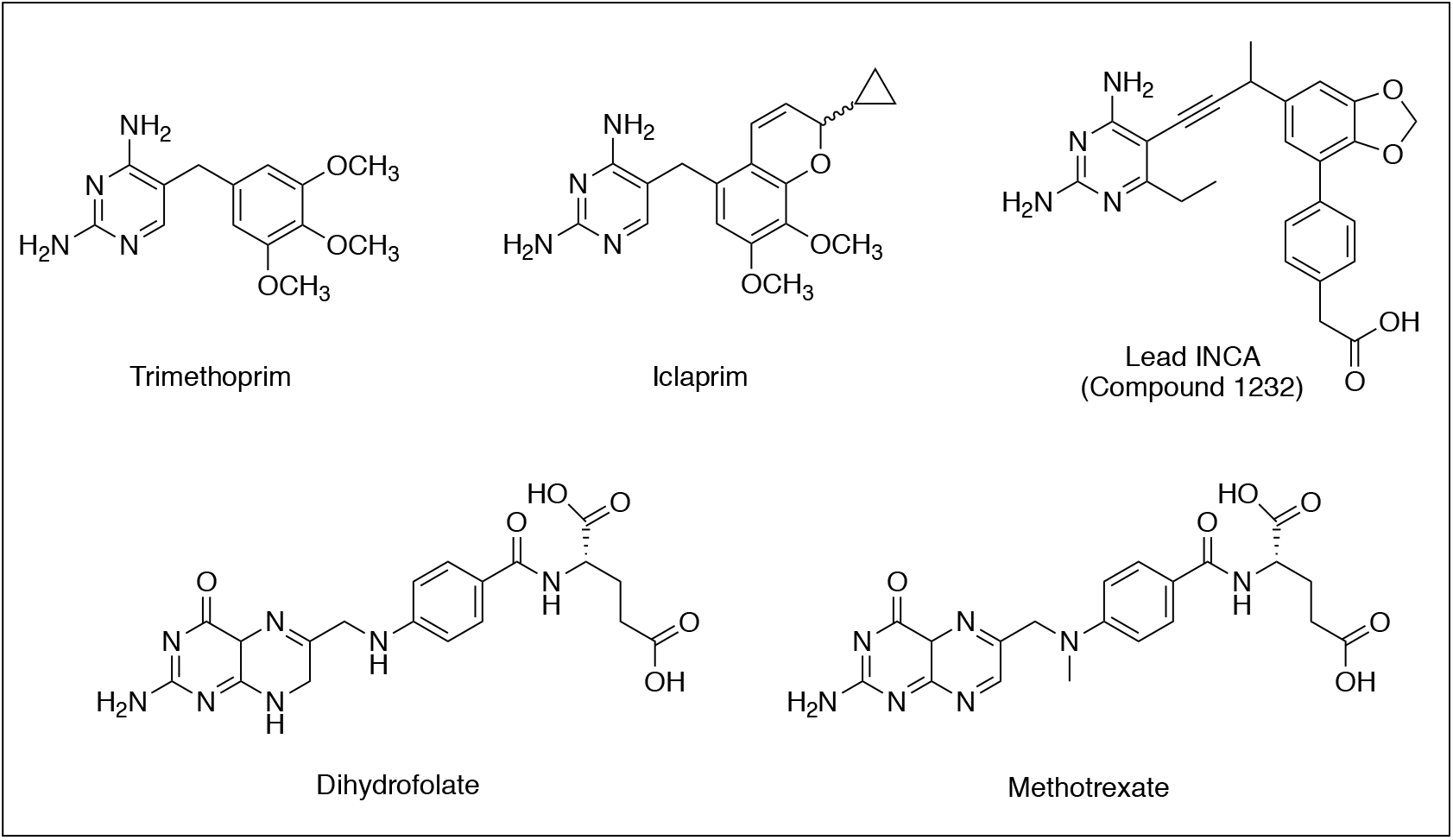
Structures of antifolates discussed in this study. Trimethoprim and iclaprim (top row) are both lipophilic antifolates with antibacterial activity. Methotrexate is a classical antifolate that mimics the natural substrate dihydrofolate. Compound **1232** is a lead ionized non-classical antifolate (INCA).

Trimethoprim resistance in *S. aureus* was first recognized in the 1980s following its clinical introduction in 1968. In the 1990s, two primary resistance mechanisms were identified as conferring clinical TMP resistance (TMP^R^): point mutations in the endogenous TMP sensitive (TMP^S^) chromosomal DHFR gene *dfrB* and the acquisition of an innately resistant DHFR enzyme, *dfrA*^5,6^. Recently two additional plasmid-encoded DHFR resistance genes, *dfrG* and *dfrK*, began appearing in MRSA infections both abroad and domestically. The *dfrG*, gene, encoding the TMP^R^ DHFR enzyme DfrG (referred to as S2 DHFR), was first isolated in Thailand and later isolated in South Africa where its import to Europe was tracked via epidemiological studies^7,8^. *dfrK*, encoding the protein DfrK, was predominately associated with agricultural, specifically swine associated infections and began recently appearing in farmers and children in farm villages in Ireland^9^. We recently identified *dfrG* and *dfrK* in clinical strains of MRSA from Connecticut hospitals, with *dfrG* being the predominant resistance determinant^10^. Our observations were mimicked in other studies identifying DfrG in as many as 78% of TMP^R^ isolates followed by *dfrA*, *dfrK* variants. Strains with mutant DfrB were seldom isolated^11,12^.

We have been developing next generation DHFR inhibitors against TMP-resistant Gram-positive^13,14^, Gram-negative^15,16^ and mycobacterial^17^ pathogens. These compounds feature a 6-ethyl-2,4-diaminopyrimidine moiety linked to a meta-biaryl system through an acetylenic linker (Figure 1). Recently, we disclosed a distinct class of antifolates designated as ionized non-classical antifolates (INCA), that are characterized by acidic functionality in the C-ring to capture the powerful interaction between the glutamate tail of classical antifolates and DHFR^14^. Importantly, this modification alters the charge distribution of INCAs to anionic/zwitterionic relative to earlier generations that are cationic/neutral. Additionally, this mono-carboxylate design allows us to exploit the key interactions used in substrate/classical antifolate binding while still maintaining the ability to passively penetrate the bacterial membrane. INCA leads exhibit strong potency against the wild-type and TMP^R^ mutant enzymes as well as clinically isolated strains containing the newly discovered *dfrG* and *dfrK* genes^10^.

With the exception of iclaprim, there has been a notable lack of development of therapeutics targeting dihydrofolate reductase in the antibacterial space. Herein, we have report a series of INCA antifolates that directly target the endogenous and acquired DHFR isoforms that confer trimethoprim and iclaprim resistant phenotypes. Using biochemical, microbiological, structural and computational techniques we are able to asses these compounds as potential antibacterial therapeutics.

## Results and Discussion

A panel of clinically isolated TMP^R^ MRSA and their corresponding DHFR enzymes, representative of the resistance landscape reported in recent literature was assembled for this study. This panel is comprised of isolates containing both a wild-type endogenous *dfrB* gene as well as either *dfrA, dfrG* or *dfrK* TMP^R^ genes, Table 1. The clinical isolates, which have been previously characterized, were collected during the course of routine clinical care from Connecticut hospitals, show unique clonality and exhibit diverse antibiotic phenotypes^10^.

**Table 1.** *Staphylococcus aureus* strains used in this study

| Strain | TMP<sup>R</sup> Determinant | Minimum Inhibitory Concentrations (µg/mL) |  |  |
| --- | --- | --- | --- | --- |
|  |  | TMP | Iclaprim | MTX |
| UCH115 | *dfrA* | 250 | 64 | >250 |
| UCH121 | *dfrG* | >1000 | >250 | >250 |
| HH1184 | *dfrK* | >1000 | >250 | >250 |
| ATCC43300 | N/A | 0.312 | 0.039 | 125 |

Of the enzymes discussed in this study, the origins, biochemical and structural features of *dfrA* have been best characterized^6,19^. DfrA has accumulated three important mutations compared to its TMP^S^ *S. epidermidis* counterpart (F98Y, G43A and V31L) that are responsible for high-level TMP resistance. While the origins of *dfrG* and *dfrK* are still unknown, it is believed that enzymes are related to *Bacillus spp.* DfrK and DfrG share a 90% sequence identity to each other, but only share a 41.5% and 42.1% sequence identity to DfrB and 38.5% and 39.8% to DfrA, respectively, Supplemental Figure S1. Despite low sequence identity, these enzymes show high homology within the active site. With the exception of a Leu5 to Ile substitution in the DfrA, DfrG, and DfrK proteins, the residues which make hydrogen bonds to the substrate, Glu27, Phe92 and Arg57, remain conserved throughout the acquired enzymes. A sequence alignment is reported in Supplemental Figure S2. DfrG and DfrK also contain the Tyr98 and Tyr149 substitutions. Mutations at these three of these positions are known to confer TMP resistance in the chromosomal DfrB enzyme^18^.

All clinical isolates used in this study exhibit high levels of antifolate resistance, Table 1. The *dfrG* and *dfrK* containing isolates, UCH121 and HH184, exhibited the highest levels of resistance to both trimethoprim and iclaprim with MIC values of >1000 μg/mL and >250 μg/mL, respectively. The *dfrA* containing strain, UCH115, also succumbs to high level antifolate resistance with MIC values of 250 μg/mL for trimethoprim and 64 μg/mL for iclaprim. Minimally, the presence of these resistant enzymes in the clinical isolates results in an 800-fold loss in cellular efficacy when compared to the TMP^S^ comparator, ATCC 43300. Overall, iclaprim is unable to evade any of these prevalent TMP resistant elements rendering the compound largely ineffective against existing TMP^R^ isolates.

In addition to cellular evaluations of *dfrG, dfrK* and *dfrA* containing strains, their corresponding recombinant enzymes, DfrG, DfrK and DfrA, were generated for kinetic and inhibitory enzymatic evaluations. Both the wild type DHFR and the TMP^R^ enzymes display the typical hyperbolic progression of Michaelis-Menten kinetics. The initial rates for DHF were applied for determination of K_M_, k_cat_ and k_cat_/K_M_ as summarized in Table 2. Kinetic analysis of K_M(DHF)_ values revealed that the TMP-resistant enzymes are comparable to the wild type DHFR. Overall, the substrate binding affinities of DfrK and DfrG are very similar to that found in DfrB, with K_M_ values of 11.01, 8.87 and 13.35 μM, respectively. DfrA displays tighter interaction with DHF with approximately a two-fold decrease in K_M_ with a value of 5.76 μM. The specificity constants (k_cat_/K_M_) of the TMP-resistant enzymes are also highly comparable to those of the wild type. A two-fold higher efficiency of DfrA enzyme, with a K_cat_/K_M_ of 0.72 μM^−1^/s^−1^, is due to the increased binding affinity to DHF while the turnover rates for the other two TMP-resistant enzymes are very similar relative to the wild type DHFR, Table 2.

**Table 2.**
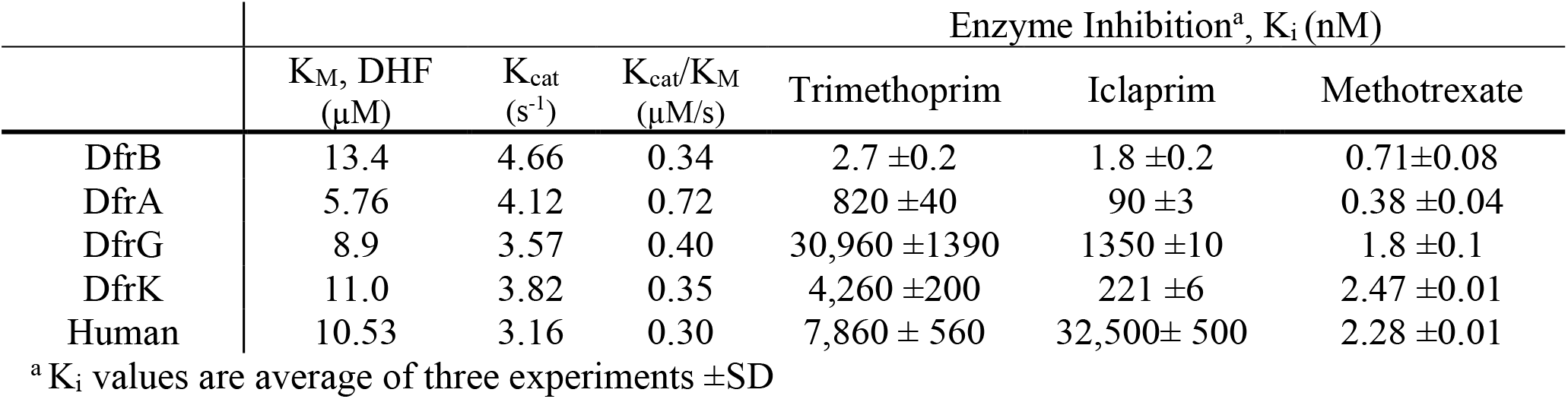
Enzyme Kinetics and Inhibition for TMP^R^ Elements and Clinical Antifolates

The resistance phenotypes observed for trimethoprim and iclaprim in the clinical isolates were recapitulated in their enzyme inhibitory activities, Table 2. DfrG conferred the highest level of resistance to both trimethoprim and iclaprim with K_i_ values of 30.96μM and 1.4μM, a >11,400 and 774-fold loss when compared to the DfrB values of 2.7 and 1.8nM respectively. Likewise, DfrA and DfrK both exhibit steep losses in affinity toward trimethoprim with K_i_ values of 820nM and 4,260nM, a >300 and >1500-fold loss potency, respectively. Iclaprim maintains higher potency against DfrK and DfrA than with DfrG with K_i_ values of 221 and 90nM, respectively. Unlike the poor inhibitory activity of the lipophilic antifolates in these enzymes, methotrexate maintains potent activity regardless of DHFR identity with K_i_ values of 0.71 nM for DfrB and 1.8, 2.47 and 0.38nM for DfrG, DfrK and DfrA, respectively.

### Design and Evaluation of Ionized Non-Classical Antifolates (INCAs)

During the last decade, we have developed and evolved the propargyl-extended antifolates from TMP like derivatives^20^ to highly functionalized inhibitors that have been tailored to the SaDHFR active site^10^. Most recently, we have developed a new class of ionized non-classical antifolates (INCAs) featuring a distal benzoic acid that added MTX-like character to the inhibitors, Figure 2. Through structure-based drug design, we have been able to establish a preliminary structure activity relationship between these benzoic acid inhibitors, interactions with Arg57 and potency. Crystal structures of the first-generation carboxylate compounds with an unsubstituted propargylic position indicated a highly coordinated water network between the *para*-benzoic acid and Arg57. Branching from the propargylic carbon with a simple methyl group displaces the biaryl system toward the Arg57 residue effectively disrupting the water network and forming one direct hydrogen bonding and one water-mediated interaction with the guanidinium side chain. While inhibitors that form water-mediated and pseudo-direct hydrogen bonding interactions with Arg57 have shown improved potency over trimethoprim, the MIC discrepancy between the DfrG, DfrK and DfrA containing strains were up to 64-fold^10^. When designing across resistant targets, it is important that the MICs across target isoforms have only small deviation to ensure the widest possible coverage. Given the broad potency of MTX against the DfrA, DfrG and DfrK enzymes, it was hypothesized that fine tuning of the interaction between the INCA carboxylate moiety and the conserved arginine sidechain would be a powerful strategy to achieving broad-based activity against these redundant DHFR containing isolates.

**Figure 2:**
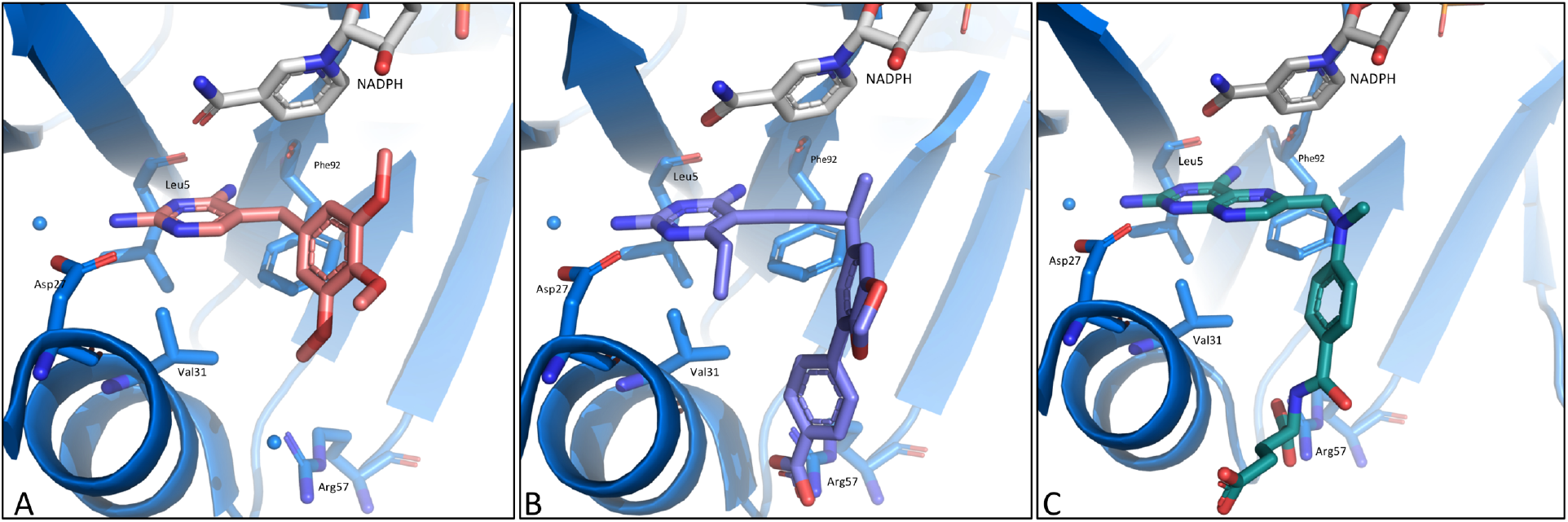
Antifolates in the SaDHFR active site A)Trimethoprim B) Compound **1191** C)Methotrexate

In order to facilitate the refinement of our INCA leads, we first obtained a crystal structure of the DfrB:NADPH:MTX ternary complex to better understand the binding mode of MTX to the bacterial reductase. In this structure, MTX makes extensive hydrogen bonding interactions with the protein’s active site including the Asp27 side chain, an active site water and the backbone carbonyls of Leu5 and Phe92. These contacts are supplemented with dual hydrogen bonds formed between the guanidinium side chain of Arg57 and the glutamate tail. The major structural difference between the human (PDB ID: 1DLS)^21^ and *S. aureus* structures is a loss of a hydrogen bonding interaction between the amide carbonyl of MTX and Asn64 side chain; this residue is replaced by a glycine in all DfrB as well as DfrA, DfrG and DfrK isoforms. Lipophilic antifolates, trimethoprim and iclaprim, maintain their interactions with the diaminopyrimidine binding pocket, however contacts with the distal Arg57 has always been an exclusive feature of classical antifolates. Moreover, the potential value of adding this functionality to antibacterial agents has been recognized as a tool to overcome resistance to point mutations, as this residue is unlikely to mutate without encountering a major fitness cost^22^.

We hypothesized that the placement of an additional carbon between the distal aryl ring and carboxylate would allow for a more productive MTX-like interaction. Therefore, a matched series of five benzoic acid and phenyl acetic acid inhibitors were synthesized, following previously reported synthetic strategies, for structural, biochemical and microbiological evaluations, Table 3.^13,16^

**Table 3.**
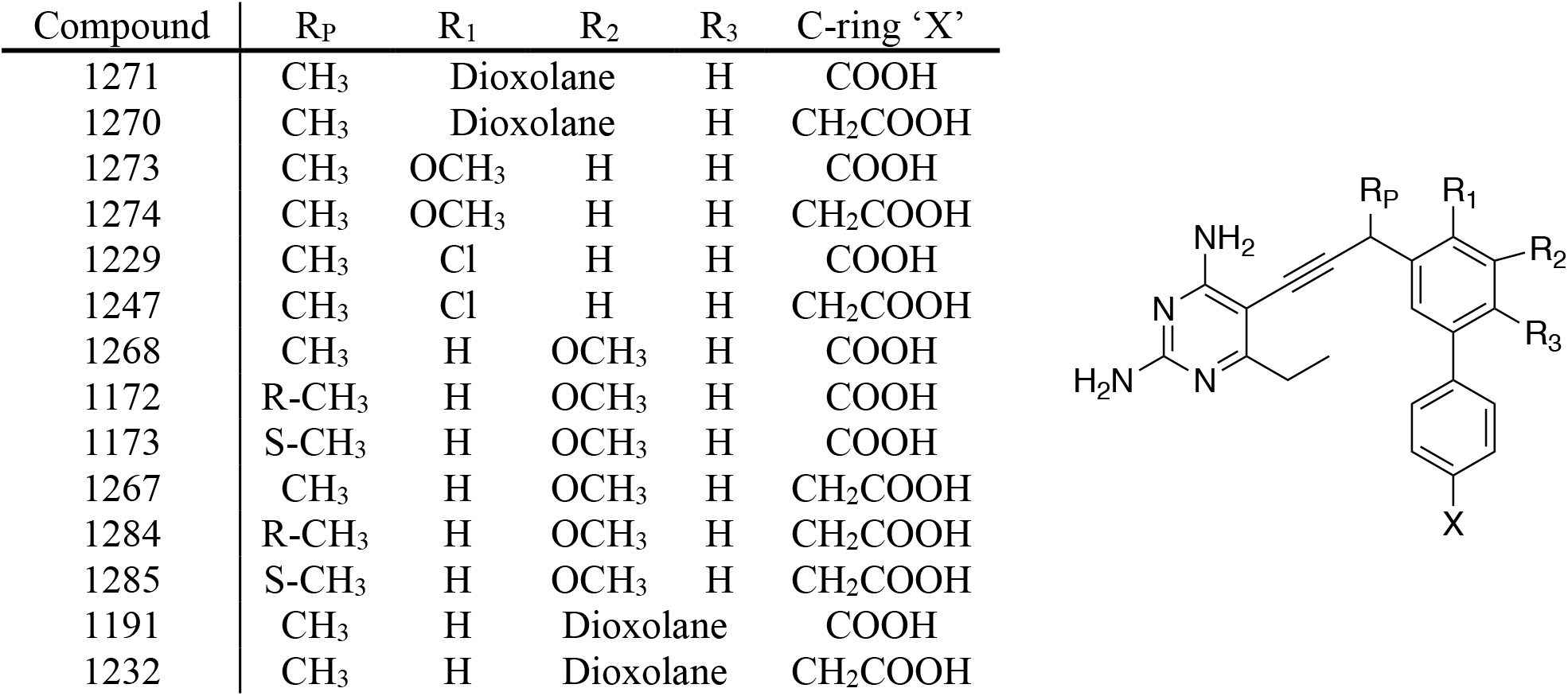
Structures of INCA Compounds

These compounds demonstrated excellent inhibitory affinity (K_i_ values <1.2 nM) toward the wild-type enzyme, DfrB. In the case of the benzoic acid series, introducing a R_1_,R_2_-dioxolane (**1271**) substituted B-ring enhanced the enzymatic inhition by 10-fold relative to a R_1_-OCH_3_ (**1273**) substituted compound. In addition, replacing the R_1_-OCH_3_ with a R_1_-Cl (**1229**) further enhanced the binding affinity by 2-fold. In comparison, the phenyl acetic acid INCAs either retained or increased their potency for DfrB relative to their benzoate congeners with R_1_-substituted B-ring systems demonstrating the most profound effects. For example, compound **1247** (R_1_-Cl) exhibited the most potent enzyme inhibition with a K_i_ of 1.2 nM, an 8-fold increase relative to its benzoic acid analog. This observation supported our hypothesis that increase in proximity and flexibility creates better interactions between the ionized extended-carboxylates and the conserved arginine. When evaluated against TMP^R^ enzymes, the INCAs were >1000-fold more potent than TMP and showed over 100-fold greater inhibitory activity relative to iclaprim against DfrA, DfrG and DfrK. Changing the B-ring substitution from R_1_-OMe to a R_1_, R_2_-dioxolane or R_1_-Cl had a positive effect across all three enzymes. Notably, **1229** showed 3, 215 and 150-fold increases in enzyme inhibition against DfrA, DfrG and DfrK, respectively, when compared to **1273**. Furthermore, all phenyl acetic acid INCAs showed enhanced inhibition against DfrA and DfrG when compared to their benzoic acid counterparts, with the most potent compound, **1267** having a K_i_ value of 2.2 nM for DfrA. For DfrK, all extended acid INCAs exhibited comparable activity to their benzoic acid partners. Based on this data, it is apparent that the extension of the anionic functionality was most beneficial for DfrA, followed by DfrG and DfrK with modest to little improvements.

Ideally, new generation DHFR inhibitors would have sufficient selectivity over human enzyme to avoid concomitant inhibition. Therefore, all new INCAs were tested against human DHFR isoform (HuDHFR) to evaluate their enzymatic selectivity. From the SAR data, it was apparent that the nature of substituents on the B- and C-rings of INCAs had an immense effect on their inhibitory activities against human DHFR. For shorter benzoic acids, moving the substitutions in B-ring from R_1_-OMe to any other position demonstrated increased selectivity, or decreased affinity towards HuDHFR. Intrestingly, extension of one carbon to the phenyl acetic acid improved the selectivity for R_1_-substituted B-ring systems, but had a detrimental effect on other B-ring systems. Notably, compound **1247** had a 33-fold increase inselectivity for the pathogenic enyzmes compared to its benzoic acid analog.

All new INCAs were evaluated for antibacterial inhibition against the panel of TMP^S^ and TMP^R^ isolates. Overall, these compounds maintained potent activity against wild-type ATCC43300 quality control strains with MIC vaues between 0.4 and <0.001 μg/mL, with the R_1_-Cl inhibitors (**1229** and **1247**) being most potent with MICs below 1 ng/mL. In general, the extension from benzoic acid to phenyl acetic acid had only a minor effect (1-2 fold increase) on potencies against the wild-type strain. Interestingly, MIC values for UCH121 and HH1184, which contain the *dfrG* and *dfrK* resistance genes, range from 0.625-10 and 0.3125 - 2.5μg/mL, respectively. The most active compounds from the benzoic acid series exhibited a >400 and >1,600-fold increase in potency compared to iclaprim and trimethoprim, respectively. For these strains, the majority of the phenyl acetic acid series were comparable or within two fold of their benzoic acid partner.

MIC values for the *dfrA* containing strain, UCH115 range from 1.25-20μg/mL, a 200 and 500-fold increase in activity when compared to trimethoprim and iclaprim. Moreover, for UCH115, the extended acids exhibited a moderate increase in potency over the benzoic acid analogs. For the R_1_,R_2_ dioxolane compounds, **1191** and **1232**, the one carbon extension reduced the MIC from 20 to 2.5μg/mL, putting the MIC within 4-fold of the DfrG and DfrK containing strains. When tested against single and double mutant strains, most of the extended acid INCAs showed a considerable increase in activity. Owing to its substantially improved activity, **1247** and **1232** have been identified as lead INCA compounds against TMP^R^ *S. aureus* pathogens.

An area of major importance in developing antibacterial DHFR inhibitors is achieving adequate selectivity over the human isoform. While the INCAs compounds tested here have less than 100-fold selectivity for the human isoform over the pathogenic enzyme, these compounds exhibit very little mammalian toxicity when tested against both MCF-10 and HepG2 cell lines. Most of the INCA compounds had IC_50_ values >200 μg/mL in both cell lines, Table 4. Compound **1229** was the most cytotoxic compound tested with IC_50_ values of 49 and 99μg/mL against MCF-10A and HepG2 cell lines, respectively, correlating with its poor enzymatic selectivity of 3.6-fold. This general lack of cytotoxicity may be attributed to the unique way in which human DHFR is regulated. It is well known that anticancer antifolates, for instance, require extraordinary target-level potency (MTX, Ki~ 5 pM)^23^ as a consequence of rapid changes to DHFR protein levels. Bastow^24^ was the first to report that MTX treatment increased the expression level of DHFR without affecting the levels of its mRNA. It was later shown that this upregulation was specific to humans^25^ and involved DHFR directly binding its cognate mRNA in the coding region^26^. Moreover, DHFR translational upregulation is an intrinsic form of resistance that protects human cells from MTX toxicity^27^. This may be the one reason that low dose MTX is well tolerated enough to allow for therapeutic applications outside of oncology. For example it is the first-line treatment for rheumatoid arthritis^28^ and is used in the management of psoriasis^29^ and ulcerative colitis^30^. We have determined that this effect is mirrored by treatment of HL-60 cells with both **1232** and MTX but not iclaprim. This indicates that MTX and INCAs induce a concentration dependent translation of human DHFR, potentially protecting the cells from the anti-HuDHFR enzymatic activity of these compounds (Supplemental Figure S3).

**Table 4.** Enzymatic inhibition of Dfr Isozymes by INCA compounds^a^ (K_i_, nM)

|  | DfrB | DfrA | DfrG | DfrK | Human Selectivity<sup>b</sup> |
| --- | --- | --- | --- | --- | --- |
| 1271 | 2.1 ±0.1 | 216 ±10 | 16 ±1 | 15 ±2 | 16.9 |
| 1270 | 2.98 ±0.09 | 9.8 ±0.9 | 13 ±1 | 6.2 ±0.2 | 23.9 |
| 1273 | 20 ±2 | 520 ±30 | 1550 ±140 | 280 ±30 | 4.4 |
| 1274 | 6.2 ±0.5 | 11.5 ±0.4 | 260 ±20 | 15.6 ±0.6 | 9.6 |
| 1229 | 10.4 ±0.2 | 15 ±2 | 7.2 ±0.5 | 1.8 ±0.2 | 2.7 |
| 1247 | 1.17 ±0.01 | 2.9 ±0.1 | 14 ±2 | 1.6 ±0.1 | 99.2 |
| 1268 | 1.76 ±0.06 | 9.7 ±0.4 | 13 ±1 | 3.4 ±0.3 | 18.9 |
| 1172 | 1.08 ±0.08 | 21.8 ±0.5 | 18 ±1 | 3.0 ±0.1 | 50.4 |
| 1173 | 1.6 ±0.1 | 15 ±1 | 16 ±1 | 9.0 ±0.8 | 46.1 |
| 1267 | 2.2 ±0.2 | 2.2 ±0.2 | 40 ±3 | 4.2 ±0.4 | 16.7 |
| 1284 | 2.1 ±0.2 | 3.0 ±0.3 | 73 ±3 | 8.1 ±0.4 | 24.7 |
| 1285 | 4.0 ±0.1 | 16 ±1 | 15 ±1 | 4.0 ±0.2 | 20.3 |
| 1191 | 1.17 ±0.02 | 17 ±1 | 6.3 ±0.4 | 4.0 ±0.6 | 74.1 |
| 1232 | 2.00 ±0.07 | 10.4 ±0.8 | 3.2 ±0.5 | 1.8 ±0.2 | 2.9 |
| MTX | 0.71 ±0.08 | 0.38 ±0.04 | 1.9 ±0.1 | 2.5 ±0.1 | 4.0 |

<sup>a</sup>K<sub>i</sub> values are average of three experiments ±SD <sup>b</sup>Human selectivity is the IC<sub>50</sub> ratio of HuDHFR to SaDHFR. Human and bacterial DHFR IC<sub>50</sub> values are reported in the Supplemental Table S1.

**Table 5.** Minimum Inhibitory Concentrations

|  | Minimum Inhibitory Concentrations (µg/mL) |  |  |  | Mammalian Toxicity IC<sub>50</sub><br>(µg/mL)<sup>a</sup> |  |
| --- | --- | --- | --- | --- | --- | --- |
|  | ATCC<br>43300 | UCH115<br>(*dfrA*) | UCH121<br>(*dfrG*) | HH1184<br>(*dfrK*) | MCF10A | HepG2 |
| 1271 | 0.005 | >10 | 0.625 | 0.312 | >200 | >200 |
| 1270 | 0.0025 | 2.5 | 1.25 | 0.625 | >200 | >200 |
| 1273 | 0.04 | 10 | 10 | 1.25 | >200 | >200 |
| 1274 | 0.04 | 5 | 10 | 1.25 | 167 ±3 | >200 |
| 1229 | <0.001 | 5 | 2.5 | 0.625 | 49.0 ± 0.7 | 99 ± 2 |
| 1247 | <0.001 | 2.5 | 2.5 | 0.312 | ND<sup>b</sup> | ND |
| 1268 | 0.004 | 10 | 1.25 | 0.625 | >200 | >200 |
| 1172 | 0.010 | 5 | 0.625 | 0.312 | 169 ±3 | >200 |
| 1173 | 0.010 | 2.5 | 5 | 2.5 | ND | ND |
| 1267 | 0.002 | 1.25 | 2.5 | 2.5 | 169 ±4 | 181 ±3 |
| 1284 | 0.010 | 1.25 | 10 | 5 | 166 ±3 | 164 ±3 |
| 1285 | 0.02 | 5 | 5 | 1.25 | >200 | >200 |
| 1191 | 0.020 | 20 | 0.625 | 0.156 | ND | ND |
| 1232 | 0.010 | 2.5 | 0.625 | 0.625 | >200 | >200 |
| MTX | 125 | >250 | >250 | >250 | ND | ND |

<sup>a</sup>Toxicity shown as the average of three independent measurements ± SD. <sup>b</sup>ND: Not determined

### Structural and Computational Studies

To aid in the understanding of the observed efficacy and to guide future optimization efforts, several crystal structures with lead compounds, **1232** and **1267**, bound to the wild type *S. aureus* DHFR were solved. Crystals of DfrB:NADPH:**1232** and DfrB:NADPH:**1267** diffracted to 1.65Å and 2.73Å, respectively. Data collection and refinement statistics are presented in Table S1. The structure of the DfrB:NADPH:**1232** complex revealed the standard five hydrogen bonding interactions between the 6-ethyl diaminopyrimidine and Asp27 side chain (2.6Å and 3.1Å), an active site water (3.0Å), Phe92 (3.1Å), and Leu5 (3.0Å) backbone carbonyls. This configuration also enables the compound to form several hydrophobic interactions between the Phe92, Leu28, Val31, Ile50 and Leu54 side chains. Additionally, the carboxylic moeity extends to form the intended dual hydrogen bonding interactions with the Arg 57 side chain, one at 2.6Å and the other at 3.1Å, Figure 3 panel B.

**Figure 3:**
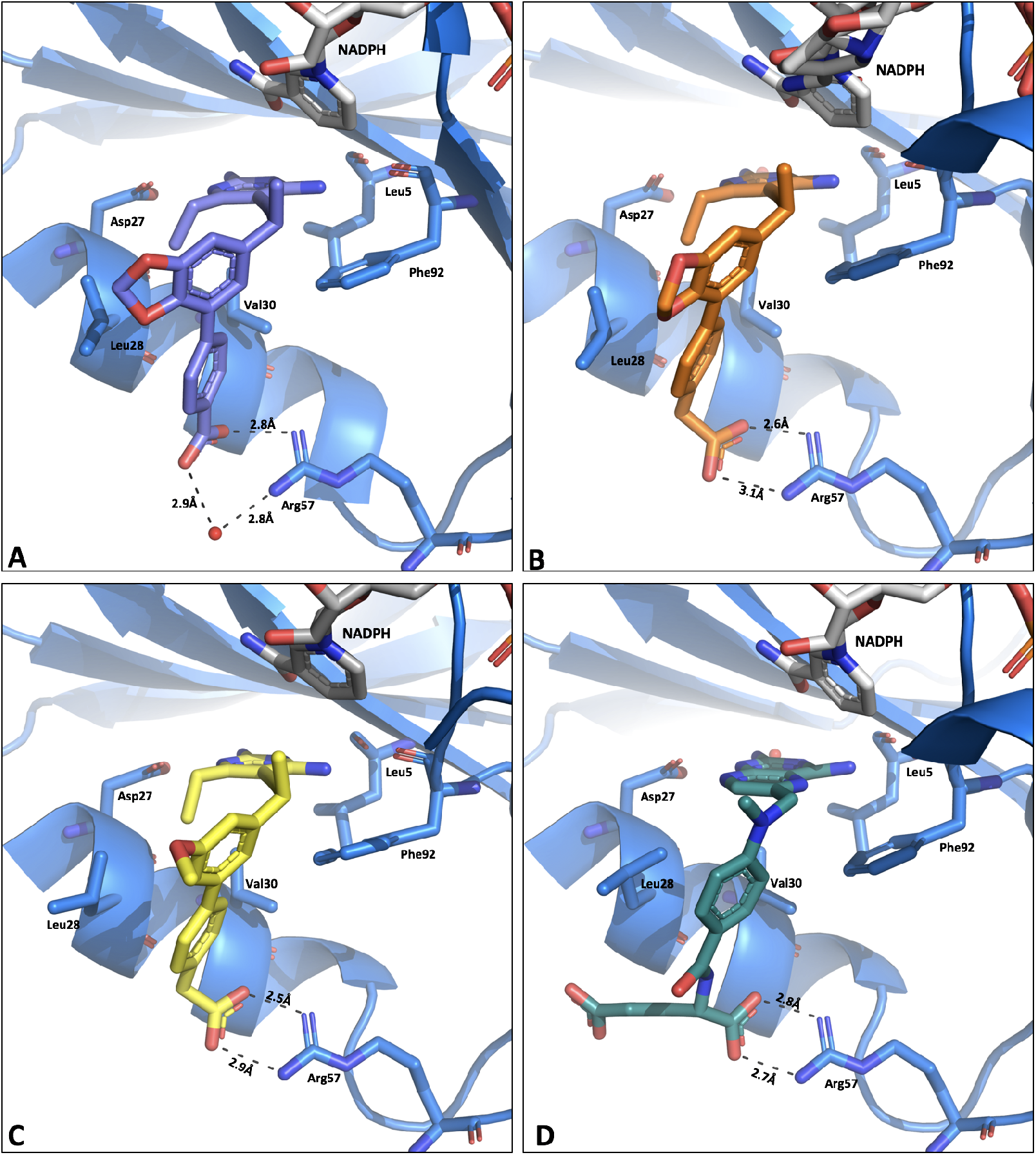
Crystal structures of A) **1191** B) **1232** C) **1267** and D) Methotrexate in the DfrB active site. This figure emphasizes the interactions between the distal acids of the INCAs/MTX and Arg57 residue. Panels B and C show the structural improvement of the phenyl acetic acid inhibitors over the benzoic acid series (panel A) and similarity to MTX (panel D).

Comparisons of its benzoic acid counterpart, **1191** (PDB ID:5JG0)^10^, reveal that the extension of the benzoic acid to phenyl acetic acid results in a 1.2Å displacement of the distal phenyl ring towards the Val31 helix. **1232** sits slightly above **1191**, 0.7Å closer to the NADPH binding pocket and this results in 2.2Å shift in the dioxolane binding. Despite the observed changes in the inhibitor binding mode, the protein’s active site appears to accommodate the altered binding positions by maintaining the rotamer orientations and hydrophobic interactions between Leu54, Leu6, Leu28 and Ile50. An overlay of these structures is presented in Supplemental Information, Figure S4.

Compounds **1232** and **1267** maintain very similar binding orientations within the *S. aureus* active site and the overall RMSD of the two structures is 0.253Å. Compound **1267** makes the conserved hydrogen bonding interactions with the active site water (2.6Å), Asp27 (2.6Å and 3.2Å), Phe92 (2.9Å) and Leu5 (2.9Å) active site residues. The most notable deviations between these two structures is the binding position of the R_2_-OMe, which extends towards the Ile 50 at a 113° angle, whereas the R_2_,R_3_-dioxolane maintains a 102° bond angle in plane with proximal aryl ring. The methoxy binding orientation results in a 1Å shift of the Ile50 helix away from the inhibitor. This movement of the adjacent helix reduces the hydrophobic interactions between the inhibitor and Ile side chain. It is believed the rigidity of the fused ring system of **1232** allows for more optimal interactions between the inhibitor and protein.

Crystals of DfrB complexed with NADPH and methotrexate diffracted to 1.80Å. This structure shows two hydrogen bonding interactions between pterin rings and the side chain of Asp27 (2.6 and 3.2Å) and backbone Leu5 (2.8Å) and Phe92 (2.9Å) as well as two hydrogen bonding interactions with Arg57 (2.7 and 2.8Å). Like in the human DHFR structures, the pterin binds in an opposite orientation than that of folate facilitating the hydrogen bonding interaction with Phe 92 (PDB ID 3FRD^31^, Supplemental Figure S5). Interestingly, in the MTX structure a rotamer of Leu28 makes more extensive hydrophobic interactions to the benzamide moiety of MTX than in the INCAs, as the binding position of the distal ring would likely clash with that residue.

In order to better understand the molecular interactions between the INCA compounds and DfrG, we constructed a homology model of DfrG active site, based on the crystal structure of DfrB bound to **1232** and NADPH. This homology model shows good overlay between the DfrB and DfrG active sites binding to **1232**, Figure 4. The DfrG structure maintains the seven hydrogen bonding interactions including Asp 27 side chain (both 3.0Å), Il5 backbone (3.1Å) and Phe92 (3.2Å) with the diaminopyrimidine and two hydrogen bonding interactions between the Arg57 and C-ring substituted phenyl acetic acid (2.9Å and 2.7Å). The hydrophobic interactions with the Val31, Ile50 and Phe92 are also maintained in these structures. Interestingly, DfrG contains a tryptophan in place of the Leu28 in the DfrB. This substitution increases the distance between the inhibitor and Leu28 (in DfrB) and Trp28 (in DfrG) from 3.6Å to 6.4Å, widening the distal region of the active site and effectively reducing the hydrophobic interactions with **1232**.

**Figure 4:**
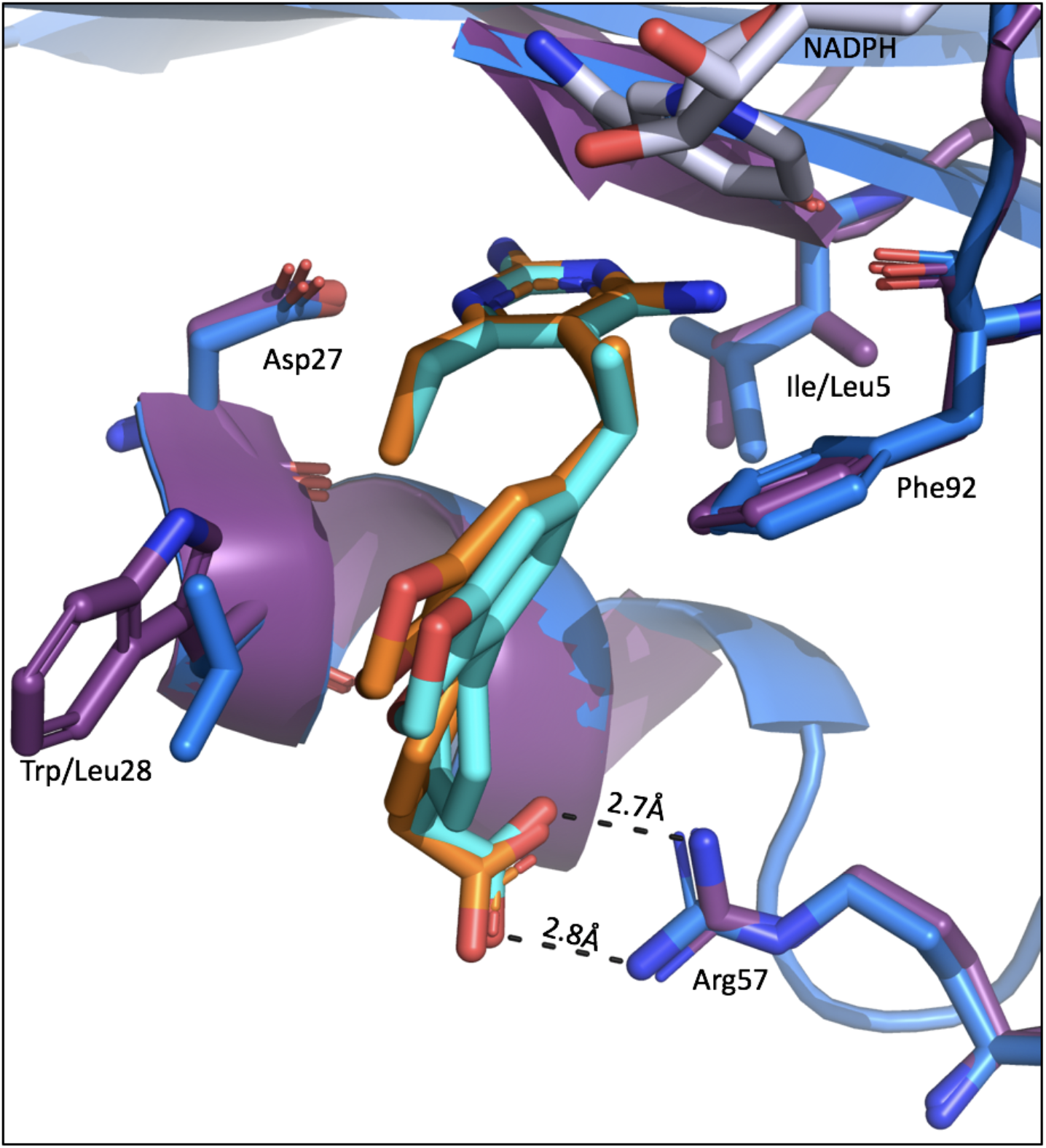
Overlay of DfrB (dark blue) with **1232**(organge) and DfrG homology model (purple) with **1232** (light blue). Active site residues shown as sticks.

Finally, we were interested in examining the conformational re-organization that these ligands undergo upon binding with bacterial DHFR and evaluating their associated energy penalties. Therefore, two different dihedrals angles, the dihedral angle between propargylic methyl and the aromatic B ring, and the dihedral angle between aromatic B and C rings were chosen for analysis. Molecular dynamic calculations performed on a paired set of benzoic acid and phenyl acetic acid ligands, **1191** and **1232** generated two separate minimum energy conformers for each ligands and were further compared with their bioactive conformers. We observed (Supplemental Table 3 and Supplemental Figure 6) that to gain the overall stabilization in the active site of the bacterial DHFR the narrow dihedral angles (4 to-18°) between propargylic methyl and aromatic B-ring were well tolerated in all of the bioactive conformers overriding the conformational bias of larger (99 to 110°) dihedral angles in the minimum energy structures. In order to make optimal interactions with the distal arginine via the dual H-bonding this dihedral angle expanded from 3.5° in shorter benzoic acid, **1191** to −16.1° in phenyl acetic acid, **1232**. Further, this overall stabilizing event also led to the 8 to 14° constrictions in the dihedral angles between the biphenyl (aromatic B and C rings) systems for phenyl acetic acid INCAs, and a 16° constriction of this dihedral angle in extended acid **1232**.

## Conclusion

Recent identification of trimethoprim resistance mechanisms in *S. aureus* has promted us to pursue the development of pan-DHFR inhibitors. Herein, we have been able to develop a hybrid class of antifolates that capture a key electrostiatic interaction common to classical antifolates without compromising the bacterial permeability associated with lipophilic antifolates. Moreover, structure-activity realtionships indicate that it should still be possible to achieve target-level selectivity even when exploiting this highly conserved interaction. It is noteworthy that developing compounds that simultaneously target both the endogenous and acquired enzymes has been a successful approach for the treatment of MRSA with 5^th^ generation cephalosporins, which unlike the earlier generation cephalosporins, target the both the endogenous PBPs as well as the acquired PBP2a.

Crystal structures of the INCA compounds disclosed here indicate several highly coordinated hydrogen bonding interactions between the inhibitor and enzyme active site, several of which have, until now, been exclusive to classical antifolates. Adding ionic functionality to earlier generations of this antifolate class revealed a highly coordinated water network between the distal region of the inhibitor and the Arg57 residue of DfrB. Lead by crystal structures and biochemical evaluations, optimization of the proximal ring system and propargylic position allowed us to displace the water network to form a single hydrogen bonding interaction to displace the water network to form a single hydrogen bond and one water mediated interaction with the active site. Now, we have identified the optimal propargylic, proximal and distal subsitutions to fully exploit the substrate binding pocket and gain potent activity across trimethoprim sensitive and resistant DHFR isoforms. Having identified several lead compounds, we can continue to improve and evaluate selectivity, determine and optimize pharmacokinetic properties and assess *in vivo* efficacy.

## Experimental

### Chemical Matter in This Study

Compounds **1172, 1173, 1191** and **1232** have been previously disclosed^10,14^. All novel compounds have been synthesized following published methods^14^. More thorough methods and compound characterization can be found in the Supplemental Information.

### Minimum Inhibitory Concentrations

Minimum inhibitory concentrations (MICs) for trimethoprim (Sigma Aldrich), iclaprim, and INCAcompounds (all in DMSO) were determined following CLSI broth dilution guidelines using isosensitest broth and an inoculum of 5×10^5^ CFU/mL^32^. MIC values were determined as the lowest concentration of inhibitor to prevent visible cell growth after 18 hour incubation at 37°C.

### Enzymatic Activity and Inhibition Assays

Enzyme activity was determined by monitoring the rate of NADPH oxidation following published methods^10,13,14^. Assays are performed at room temperature in a buffer solution 20 mM TES, pH 7.5, 50 mM KCl, 0.5 mM EDTA, 10 mM beta-mercaptoethanol and 1mg/mL BSA. For enzymatic assays (500 μL volume reactions), 1 μg of protein is mixed with 100 μM of NADPH and the reaction is activated with 100μM DHF (in 50 mM TES, pH 7.0). The reaction is monitored in a spectrophotometer at A_340_. The steady-state kinetic parameters of K_M(DHF)_ and K_i DHF_ were obtained for TMP^R^ enzymes and compared to the wild type DHFR. Michaelis-Menten constants (K_M_ and V_max_) were graphically determined for the substrate from the initial rates at various DHF concentration (1.6 to 100 μM) and NADPH saturation (100 μM), using a non-linear least-squares fitting procedure^33^. The turnover number (*k*_cat_) was calculated on the basis of the enzyme molecular mass. K_i DHF_ values were obtained using Cheng-Prusoff equation^34^. The reported data are averages of two independent experiments, where each experiment was conducted in triplicates.

For enzyme inhibition experiments, 1 μg of protein is mixed with 100μM of NADPH and varying concentrations of inhibitor for 5 minutes. After 5 minutes, the reaction is activated with 100μM DHF (in 50mM TES, pH 7.0) and monitored at A340. The IC_50_ is defined as the concentration of compound required to reduce the activity of protein by 50%. For comparisons across Dfr species, the IC_50_ values are converted to K_i_ to account for differing substrate affinities

### HepG2 and MCF-10 Cytotoxicity

Adherent cell lines were maintained in Eagle’s Minimal Essential Media with 2 mM glutamine and Earle’s Balanced Salt Solution adjusted to contain 1.5 g/L sodium bicarbonate, 0.1 mM non-essential amino acids, 1 mM sodium pyruvate and 10 % fetal calf serum. Fetal calf serum used in these assays was lot matched throughout. All cultures were maintained under a humidified 5 % CO2 atmosphere at 37°C, had media refreshed twice weekly and were subcultured by trypsinization and resuspension at a ratio of 1:5 each week. Toxicity assays were conducted between passages 10 – 20. Target compound toxicity was measured by incubating the test compound with the cells for four hours, washing the cells and finally treating the cells with Alamar Blue. After 12 – 24 hours the fluorescence of the reduced dye was measured. Fluorescence intensity as a function of test compound concentration was fit to the Fermi equation to estimate IC_50_ values.

### Protein Preparation

Purification for all proteins in this study have been previously published^10,13,14,35^. In brief, proteins were expressed in BL21(DE3) E. coli cells with 1mM IPTG induction and 18 hour post induction growth at 18°C. Cells were lysed via sonication in a buffer of 25 mM Tris, pH 8.0, 0.4 M KCl supplemented with 0.1 mg/mL lysozyme, DNase, RNase and a cOmplete Mini Protease Inhibitor tablet. Enzymes were purified using Ni-NTA chromatography washing the bound protein with a solution of 25 mM Tris, pH 8.0 and 0.4 M KCl. Protein was eluted with 25mM Tris, pH 8.0, 0.3 M KCl, 20% glycerol, 0.1 mM EDTA, 5 mM DTT and 250 mM imidazole. Elution fractions were run on SDS-PAGE gel and pure protein was pooled and desalted into a buffer of 25 mM Tris, pH 8.0, 0.1 M KCl, 0.1 mM EDTA and 2 mM DTT and flashed frozen for storage at −80°C.

### Protein Crystallography

#### DfrB:NADPH:MTX

DfrB at 13 mg/mL was incubated with 1 mM of MTX (in DMSO, Sigma Aldrich) and 2 mM of NADPH (in water, Sigma Aldrich) for several hours. The solution was pelleted at 4°C to remove any insoluble or precipitated protein. The protein was crystallized at 4°C in a 1:1 ratio in a solution of 0.1 M MES, pH 5.5, 0.2 M sodium acetate, 15% PEG 10,000 (Hampton Research) and 20% gamma-butyrolactone (Sigma Aldrich) as an additive. Crystals generally grew within 7 days and were flash frozen in solution containing 25% glycerol.

#### DfrB:NADPH:**1232**

DfrB at 13mg/mL was incubated with 1mM of **1232** (in DMSO) and 2mM of NADPH (in water) for several hours. The solution was pelleted at 4°C to remove any insoluble or precipitated protein. The protein was crystallized at 4°C in a 1:1 ratio in a solution of 0.1 MES, pH 6.0, 0.1M sodium acetate, 15% PEG 10K and 20% gamma-butyrolactone as an additive. Crystals generally grew within 7 days and were flash frozen in solution containing 25% glycerol.

#### DfrB:NADPH:**1267**

DfrB at 13mg/mL was incubated with 1mM of **1267** (in DMSO) and 2mM of NADPH (in water) for several hours. The solution was pelleted at 4°C to remove any insoluble or precipitated protein. The protein was crystallized at 4°C in a 1:1 ratio in a solution of 0.1 MES, pH 6.0, 0.3M sodium acetate, 17% PEG 10,000 and 20% gamma-butyrolactone as an additive. Crystals generally grew within 7 days and were flash frozen in solution containing 25% glycerol.

All data were collected at Stanford Synchrotron Radiation Light (SSRL), SLAC National Accelerator Laboratory. Data were indexed using HKL2000. Phaser was used to identify molecular replacement solutions using PDB ID: 3F0Q^35–38^.

### DfrG:NADPH:**1232** Homology Modelling

Homology modeling of DfrG active site was accomplished via the study of extant DHFR crystal structures in complex with various ligands. In this case, the DfrB:NADPH:**1232** crystal structure was selected as the input starting structure for the homology modeling of DfrG active sites. Next, an intermediate model was generated using a structure prediction calculation, termed “OSPREY-designed sequence replacement” (ODSR). This process involves mutation to the target sequence implemented by side chain replacement. Here, all residues within 8Å of **1232** were selected and mutated to the appropriate DfrG amino acid, determined by sequence alignment to the sequence of DfrG. Sequence alignment was performed using CLUSTAL X 2.1 software^39^. Subsequently, side-chain replacement and global minimum energy conformation (GMEC) calculation were performed using OSPREY^40,41^. Following ODSR, the intermediate model was all-atom minimized using the SANDER package from the AMBER biomolecular simulation package^42^. Minimization was allowed to proceed for 1,000 steps, resulting in a fully-minimized homology model for DfrG active sites in complex with **1232** and NADPH. Scripts are available upon request for all steps in our protocol.

## Ancillary Information

## Supporting information

File S1: DfrG active site homology model with **1232** (PDB Format)

Figure S1: Sequence similarity comparison of DHFR enzymes used in this study

Figure S2: Sequence alignment of DHFR enzymes used in this study

Figure S3 DHFR expression in response to antifolate exposure

Figure S4: Overlay of **1191** and **1232** with active site amino acids

Figure S5: Overlay of folate and methotrexate in the DfrB active site

Figure S6: Minimum energy structures using biphenyl and propargylic dihedral drive

Table S1: DfrB and Human DHFR IC_50_ Values used to Calculate Human Selectivity

Table S2: Crystallography Data Collection and Refinement Statistics

Table S3: Comparison of dihedral angles in minimum energy and bioactive conformations Supplemental Biological and Synthetic Methods and Compound Characterization

Figures S7-S12: ^1^H NMR Spectra of Novel Compounds

## PDB Codes

DfrB:NADPH:**1232 | PDB ID: TBD**

DfrB:NADPH:**1267 | PDB ID: TBD**

DfrB:NADPH:Methotrexate | **PDB ID: TBD**

## Acknowledgements

We acknowledge funding from the National Institutes of Health grants GM078031 and GM078031 to BRD and AI111957 and AI104841 to DLW. We also acknowledge the SSRL beamline staff for their assistance in remote data collection and crystallography support. Use of the Stanfrod Synchrotron Radiation Lightsource, SLAC National Accelerator Laboratory, is supported by the U.S. Department of Energy, Office of Science, Office of Basic Energy Sciences under Contract No. DE-AC02-76SF00515.

## Author contributions

SMR: Strain characterization and susceptibilites, enzyme inhibition, protein purification and protein crystallography. DS, YY and KV: synthesis and characterization of novel chemical matter. JK: protein purification and enzyme kinetic experiments. SW, GTH and MSF: DfrG homology models. SS: DHFR expression profiling. NDG and JBA: preformed cytotoxicity assays. NDP: oversaw cytotoxicity experiments. AW: oversaw DHFR expression and western blotting experiments. BRD: oversaw computational experiments and DLW directed biology and synthetic chemistry. SMR, DJ, and DLW wrote and SMR, DJ, JK, DLW edited the manuscript.

## Abbreviations

IC_50_: concentration for 50% inhibitory activity
MIC: Minimum inhibitory concentrations
TMP^R^: Trimethoprim resistant
DHFR: dihydrofolate reductase

